# Quantifying acyl-chain diversity in isobaric compound lipids containing monomethyl branched-chain fatty acids

**DOI:** 10.1101/2024.05.28.596332

**Authors:** CR Green, MJ Kolar, GH McGregor, AT Nelson, M Wallace, CM Metallo

## Abstract

Compound lipids comprise a diverse group of metabolites present in living systems, and metabolic- and environmentally-driven structural distinctions across this family is increasingly linked to biological function. However, methods for deconvoluting these often isobaric lipid species are lacking or require specialized instrumentation. Notably, acyl-chain diversity within cells may be influenced by nutritional states, metabolic dysregulation, or genetic alterations. Therefore, a reliable, validated method of quantifying structurally similar even-, odd-, and branched-chain acyl groups within intact compound lipids will be invaluable for gaining molecular insights into their biological functions. Here we demonstrate the chromatographic resolution of isobaric lipids containing distinct combinations of straight-chain and branched-chain acyl groups via ultra-high-pressure liquid chromatography (UHPLC)-mass spectrometry (MS) using a C30 liquid chromatography column. Using metabolically-engineered adipocytes lacking branched-keto acid dehydrogenase A (Bckdha), we validate this approach through a combination of fatty acid supplementation and metabolic tracing using monomethyl branched-chain fatty acids and valine. We observe resolution of numerous isobaric triacylglycerols and other compound lipids, demonstrating the resolving utility of this method. This approach strengthens our ability to quantify and characterize the inherent diversity of acyl chains across the lipidome.

## Introduction

Monomethyl branched-chain fatty acids (BCFAs) are a group of bioactive lipids garnering increased attention for their potential significance in health and disease(1–4). Monomethyl BCFAs contain an additional methyl group on the penultimate (*iso*) or antepenultimate (*anteiso*) carbon of a fatty acid (Fig. 1A). They are synthesized *de novo* in mammals(5), worms(6), and microbes(7) from branched-chain amino acid (BCAA) catabolic intermediates. BCFAs are altered in the contexts of metabolic syndrome and obesity/weight-loss(8–10) and the abundance of odd-chain fatty acids (OCFAs) and BCFAs is influenced by cobalamin (vitamin B12) deficiency(11), suggesting such measurements can provide insights into disease or nutritional status. Additionally, BCFAs are abundant in certain sources of dietary fat, including pasture-raised eggs(12), fish(13), and animal or dairy products(14). Currently, there is limited data on esterified BCFAs in animal and human tissue in part because of their difficulty to measure on platforms that rely on liquid chromatography (LC). The composition of compound lipids like triacylglycerols often include acyl chains derived from endogenous and dietary sources. Characterizing the precise makeup of these lipids within biological systems poses significant challenges owing to their vast diversity. Moreover, the structural isomerism of BCFAs in comparison to straight-chain fatty acids further compounds the challenges associated with chromatographic separation.

**Figure 1.**
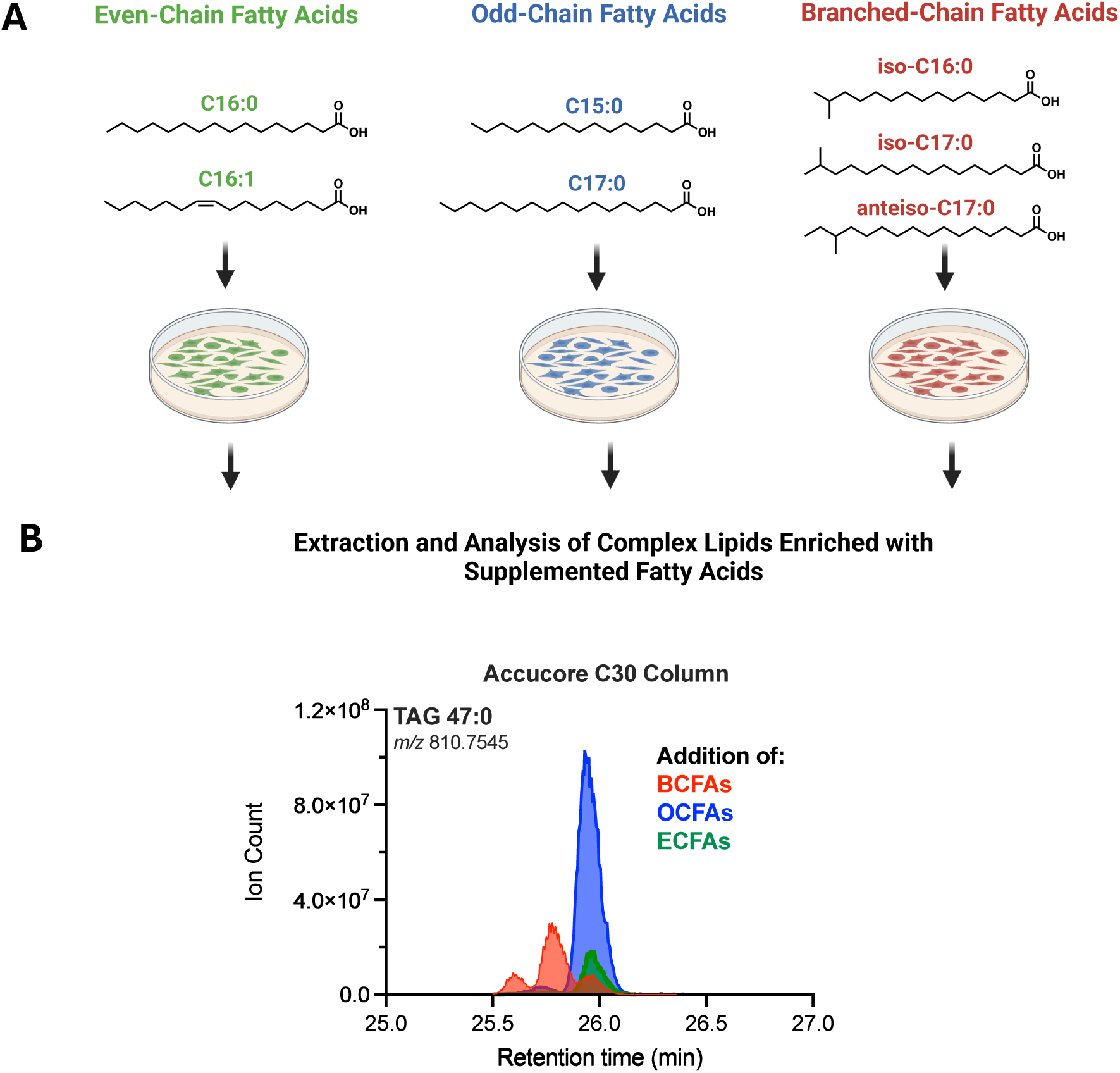
Method development for the chromatographic resolution of isobaric triacylglycerols. (**A**) Structures of even-chain-, odd-chain-, and branched-chain fatty acids and schematic illustrating the respective lipid mixtures added to 3T3-L1 adipocytes for analysis of their incorporation in complex lipid structures. The following lipid mixtures (100 µM per lipid) and ratios were used for each condition: 1:1 palmitic acid (C16:0): palmitoleic acid (C16:1), 1:1 pentadecanoic acid (C15:0): heptadecanoic acid (C17:0), 1:1:1 iso-C16:0:iso-C17:0: anteiso-C17:0. (**B**) Overlay of extracted ion chromatograms of TAG 47:0 (m/z 810.7545 ± 5 ppm) from each experimental condition supplementing 3T3-L1 adipocytes with either ECFAs (green), OCFAs (blue), or BCFAs (red) using UHPLC-MS equipped with a and Accucore C30 (2.6 µm, 250 x 2.1 mm, Thermo). Chromatographic resolution was also assessed using a C18 column (**Supplemental Fig. 2**).

BCFAs are resolvable from their isomeric straight-chain counterparts as fatty acid methyl esters (FAMEs) via gas chromatography. However, separation on LC platforms of compound lipids containing different isobaric acyl chains is more challenging. Furthermore, the lack of readily available standards has made validation challenging. Recent studies have successfully employed C30 reverse phase chromatography to separate ethyl-branched fatty acids(15), cis/trans isomers and sn-positions of lysophospholipids(16) intra-class lipid isomers and head group modifications(17), and triacylglycerol isomers(18, 19). Therefore, we examined the ability of this column to separate monomethyl BCFA-containing compound lipids produced within biological materials.

Here we demonstrate chromatographic resolution of BCFAs in compound lipids using an Accucore C30 column and ultra-high-pressure liquid chromatography (UHPLC) coupled to high-resolution orbitrap mass spectrometry. The typical workflow for identifying novel lipids entails synthesizing standards, a process that is both labor-intensive and costly, especially when a synthetic route is not readily available. To circumvent this, we validate our findings using metabolically-engineered 3T3-L1 adipocyte cultures and stable isotope tracing using ^2^H-iso-C16:0 or ^13^C-valine. This approach successfully resolved isobaric, structurally distinct acyl-chains within triacylglycerols (TAGs), phosphatidylcholine (PC), phosphatidylethanolamine (PE), ceramide (Cer), and sphingomyelin (SM) in biological samples. Overall, this approach offers a means to validate novel lipid species without relying on synthetic standards and expands our ability to identify isomeric acyl-chain distributions across the lipidome. Given the complexity of BCFA synthesis in mammals, such approaches may provide insight on tissue-specific lipogenesis, BCAA catabolism, and the molecular impacts of vitamin B12 deficiency.

## Results

### Chromatographic separation of BCFA-containing TAGs using a C30 column

As an initial point of comparison, we and others typically quantify BCFAs as saponified fatty acid methyl esters via GC-MS(20, 21). For example, *n*- (straight chain), *iso*-, and *anteiso*-fatty acids are readily separated using a mid-polarity DB-35MS column from complex lipid mixtures such as lanolin or 3T3-L1 adipocytes, biological samples with a high abundance of BCFAs(5, 22) (Supplemental Fig. S1). 3T3-L1 adipocytes increase BCAA catabolism and BCFA synthesis during differentiation(5, 10). To drive the biosynthesis of triacylglycerols containing even-chain fatty acids (ECFAs), odd-chain fatty acids (OCFAs) and BCFAs we treated 3T3-L1 adipocytes with the following BSA-complexed mixtures at 100 µM for 96 hours: a) 1:1 C16:0: C16:1, b) 1:1 C15:0: C17:0, or c) 1:1:1 iso-C16:0: iso-C17:0: anteiso-C17:0, respectively (Figure 1A).

We analyzed each extract on a Thermo Q-Exactive orbitrap-MS coupled with a UHPLC equipped with either a C18 column (Supplemental Fig. S2) or an Accucore C30 column (Fig. 1B). We focused on TAG 47:0 (*m/z* 810.7545 corresponding to the [M+NH4]^+^ adduct ion) as a principal representation due to its notable abundance and the presence of different peak retention times, suggesting the potential incorporation of the various fatty acids supplemented into the culture medium. There are numerous other compound lipids where this exercise may be performed. The C18 column resolved this species as one peak which was slightly shifted in BCFA-treated culture extracts (Supplemental Fig. S2). In contrast, TAG 47:0 in BCFA-treated cultures resolved as three peaks on C30 chromatography (red), while both arms containing straight-chain fatty acids (blue and green) resolved TAG 47:0 as a single, overlapping peak (Fig. 1B). These data indicate that C30 chromatography effectively separates TAGs containing distinct BCFA isomers but not TAGs containing combinations of straight-chain acyl groups of different length. Of note, odd-chain fatty acids (OCFAs) are high in differentiated 3T3-L1 adipocytes when cultured in typical DMEM +10% FBS due to vitamin B12 deficiency (Fig. 1a-b, blue peaks)(23, 24).

We next employed MS2 analysis to elucidate the acyl composition of TAG 47:0. For instance, by tracking the neutral loss of NH_3_ and an acyl side chain from the original TAG to a diacyl product ion, we can determine the mass of the lost acyl group (Table 1). In analyzing the peak where ECFAs were supplemented, the predominant MS2 ions are m/z 537.4864 and 551.5020 at a 2:1 ratio, suggesting loss of C16:0 or C15:0 acyl chains (Fig. 2, Table 1). These data suggest that the predominant TAG 47:0 peak has two acyl chains containing C16:0 and one C15:0. Additionally, noteworthy are the less abundant MS2 ions with m/z 523.4704 and 565.5155, indicating loss of C17:0 and C14:0, respectively. This suggests the presence of acyl chains with such composition within this peak as well. The treatment of adipocytes with OCFAs (blue) overlapped with the control extract, indicating a comparable composition of acyl chains but with higher overall abundance. We also detected lower abundances of TAG 47:0 species MS2 fragments with the loss of C14:0, C17:0, and C18:0 (Fig. 2).

**Figure 2.**
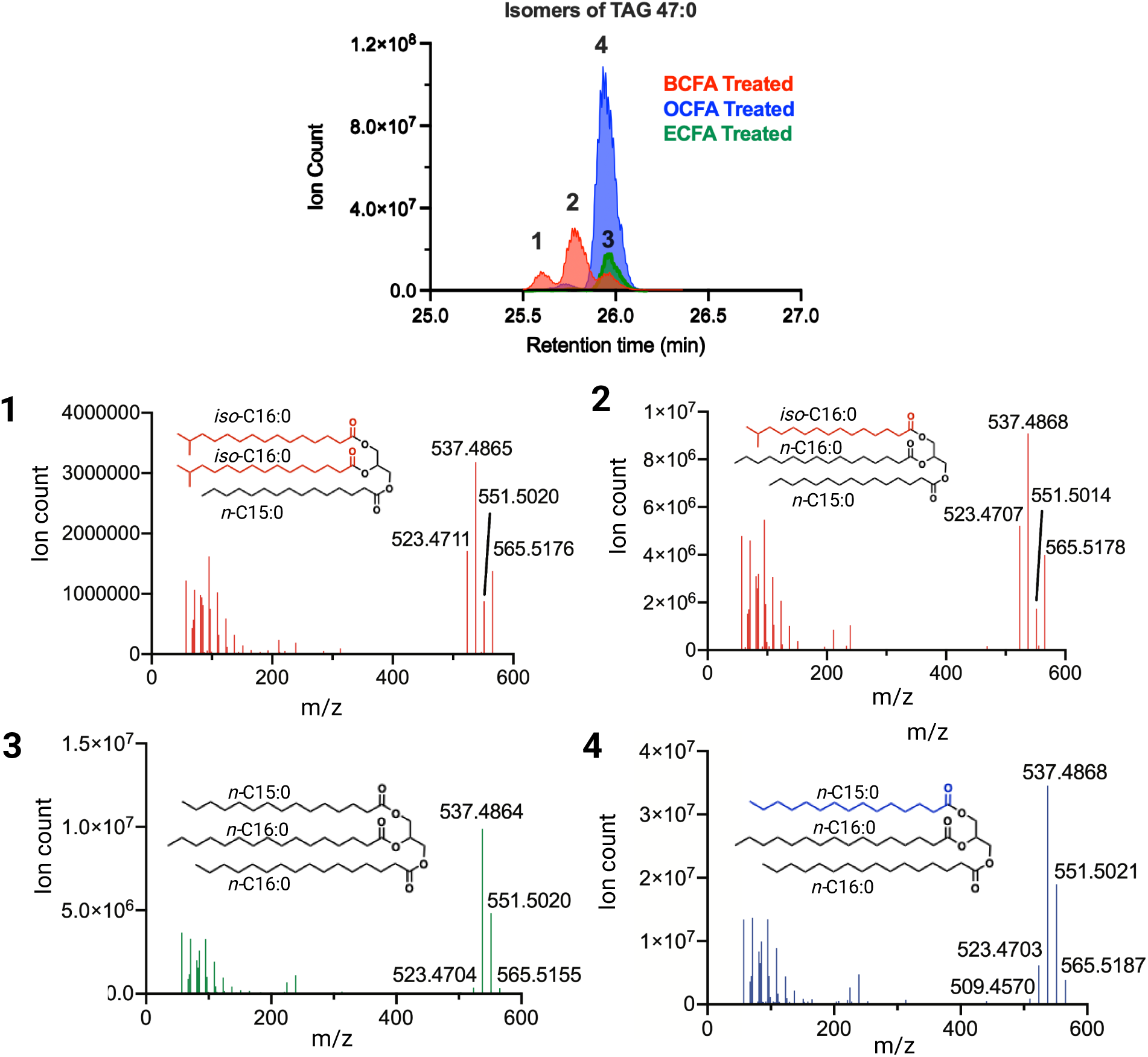
Identification of BCFA-containing TAGs. Overlay of extracted ion chromatograms of TAG 47:0 [M+NH_4_]^+^ ion in each FFA treated condition with respective MS2 spectra. Subpanel **1-4**, showing the MS2 spectra of each analyzed peak (1–4). A theoretical structure of the most abundant TAG 47:0 per MS2 spectra is shown. Note: we are unable to determine the stereospecific acylation patterns from MS2 spectra.

**Table 1.**
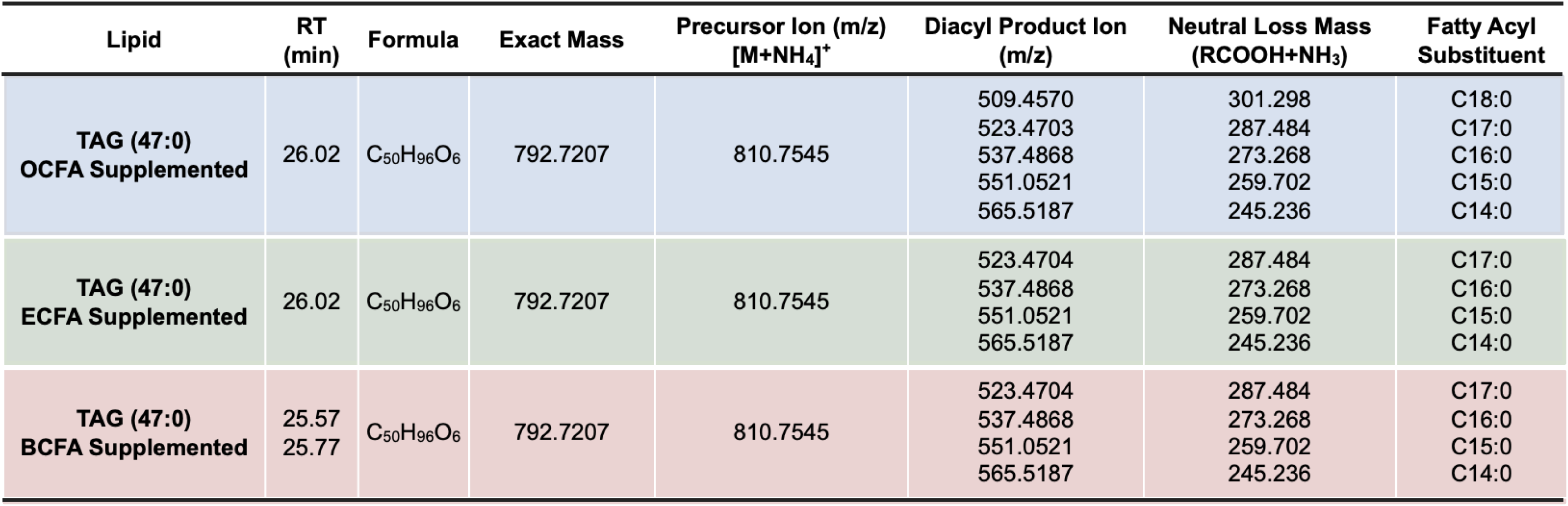
Table summarizing the acyl chain composition of the [M+NH_4_]^+^ TAG 47:0 ion via MS2 in each of the treated conditions. The TAG [M+NH_4_]^+^ precursor ion is subjected to collision-induced-dissociation (CID) and undergoes the neutral loss of a fatty acyl group and ammonia (RCOOH + NH_3_) resulting in a diacylglycerol (DAG) fragment ion. Subtraction of ammonia from the neutral loss mass provides the exact mass of the fatty acyl substituent that was lost.

| Lipid | RT (min) | Formula | Exact Mass | Precursor Ion (m/z)<br>[M+NH<sub>4</sub>]<sup>+</sup> | Diacyl Product Ion (m/z) | Neutral Loss Mass (RCOOH+NH<sub>3</sub>) | Fatty Acyl Substituent |
| --- | --- | --- | --- | --- | --- | --- | --- |
| **TAG (47:0)**<br>**OCFA Supplemented** | 26.02 | C<sub>50</sub>H<sub>96</sub>O<sub>6</sub> | 792.7207 | 810.7545 | 509.4570<br>523.4703<br>537.4868<br>551.0521<br>565.5187 | 301.298<br>287.484<br>273.268<br>259.702<br>245.236 | C18:0<br>C17:0<br>C16:0<br>C15:0<br>C14:0 |
| **TAG (47:0)**<br>**ECFA Supplemented** | 26.02 | C<sub>50</sub>H<sub>96</sub>O<sub>6</sub> | 792.7207 | 810.7545 | 523.4704<br>537.4868<br>551.0521<br>565.5187 | 287.484<br>273.268<br>259.702<br>245.236 | C17:0<br>C16:0<br>C15:0<br>C14:0 |
| **TAG (47:0)**<br>**BCFA Supplemented** | 25.57<br>25.77 | C<sub>50</sub>H<sub>96</sub>O<sub>6</sub> | 792.7207 | 810.7545 | 523.4704<br>537.4868<br>551.0521<br>565.5187 | 287.484<br>273.268<br>259.702<br>245.236 | C17:0<br>C16:0<br>C15:0<br>C14:0 |

In the TAG 47:0 chromatogram from BCFA-treated adipocytes (red), three notable peaks are detected, unlike the one peak that was seen with the ECFA and OCFA supplemented cells. One peak overlaps with the prior conditions, and two peaks elute earlier at 25.77 and 25.57 min. Given the emergence of these two peaks exclusively in the BCFA-treated adipocytes, it is reasonable to infer that the supplemented BCFAs are incorporated into the TAG 47:0. This inference is supported from the earlier elution pattern indicating decreased hydrophobicity which is likely due to BCFA incorporation, as free BCFAs are less hydrophobic compared to their straight-chain counterpart(25). The MS2 ions of these peaks revealed the loss of the following acyl chains in decreasing abundance C16:0 > C17:0 > C14:0 > and C15:0. The earlier retention time of these C16:0- and C17:0-containing TAG 47:0 suggests the incorporation of supplemented BCFAs (Table 1).

### 2H7-iso-C16:0 tracing reveals BCFA incorporation across lipid classes

To further validate the incorporation of BCFAs into TAGs, we engineered 3T3-L1 adipocytes using CRISPR-Cas and sgRNA targeting branched-keto acid dehydrogenase A (*Bckdha*) and performed subsequent metabolic tracing with ^2^H_7_-iso-C16:0. Bckdha mediates the rate-limiting step of BCAA catabolism, and knockdown of this enzyme prevents the *de novo* biosynthesis of BCFAs. The addition of isotopically labeled iso-C16:0 BCFA to these Bckdha-KO adipocytes reliably confirms the structure of the observed peaks. We infected pre-adipocyte cultures with lentivirus delivering CRISPR-Cas and sgRNAs, confirming loss of Bckdha and BCAA catabolism after differentiation(26). These cells were treated with 100 mM ^2^H_7_-iso-C16:0-BSA complex for 96 hours followed by lipid extraction and UHPLC-MS analysis. For simplicity, we focused on TAG 48:0 because complete incorporation of all the acyl chains with our deuterated iso-C16:0 standard would give us a TAG 48:0 species. The TAG 48:0 species without incorporation of ^2^H_7_-iso-C16:0, eluted at 26.32 min (m/z 824.7702) (Fig. 3A, black peak). Notably, these acyl chains are presumed to all be straight chain considering the absence of BCFA production in the Bckdha-KO adipocyte model(26). Next, we examined TAG 48:0 species containing a tracer-induced mass shift due to ^2^H_7_-iso-C16:0 incorporation (Fig. 3A). Compared to the TAG 48:0 with all straight chain C16:0 acyl chains, we observed three distinct peaks with earlier elution times which correspond to TAG 48:0 species with single, double, and triple incorporated ^2^H_7_-iso-C16:0 acyl-chains. We identified a peak at 26.09 min (peak 3)which contained MS2s in a 2:1 ratio of m/z 558.5460 and 551.5019, indicating the presence of a single ^2^H-iso-C16:0 acyl-chain (Table 2). Similarly, a TAG 48:0 molecule containing two ^2^H_7_-iso-C16:0 acyl chains (14 ^2^H isotopes) eluted at 25.89 min (peak 2) and had a 2:1 ratio of 558.5460 and 551.5019 MS2 fragments (Table 2). Finally, we also detected TAG 48:0 species entirely comprised of BCFAs that contained 3 ^2^H_7_-iso-C16:0 tracer molecules (21 ^2^H isotopes), eluting at 25.66 (peak 1) min (Fig. 3A, Table 2). These samples were also analyzed on a C18 column, but we were unable to achieve suitable separation (data not shown).

**Figure 3.**
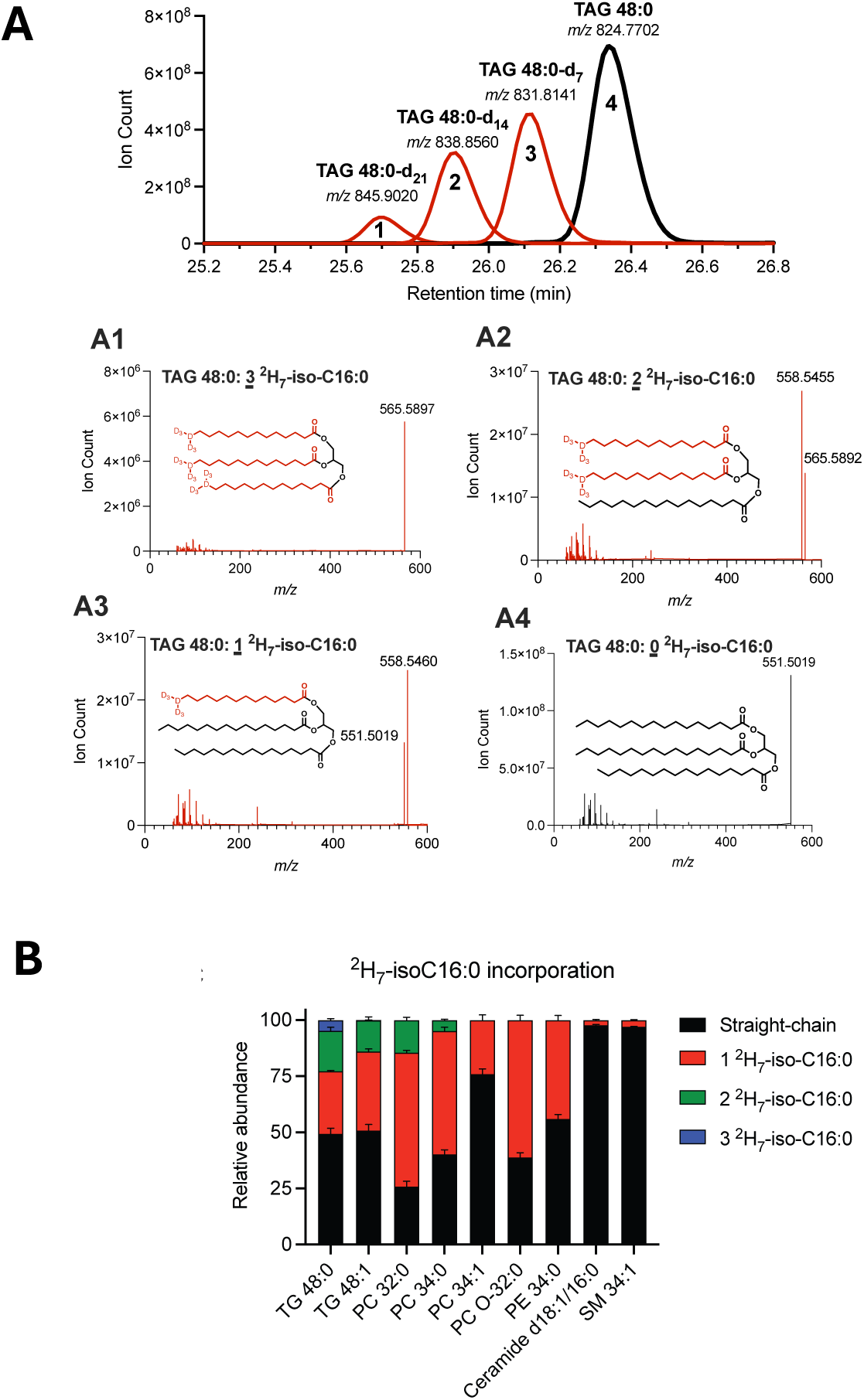
^2^H_7_-iso-C16:0 tracing confirms BCFA incorporation into TAGs and other compound lipids. **(A)** Overlay of extracted ion chromatograms of TAG 48:0 (control), TAG-48:0-d_7_, TAG-48:0-d_14_ and TAG 48:0-d_21_ representing the incorporation of 0, 1, 2, or 3 ^2^H_7_-iso-C16:0 acyl chains respectively. Bckdha-KO adipocytes cultured with 500 nM of cobalamin were treated with 100 µM ^2^H_7_-iso-C16:0. Subpanel **A1-A4** showing the respective MS2 spectra and structure of each peak. Deuterium represented by d, D, or ^2^H. **(B)** Stacked plot depicting the percentage of all straight-chain-containing or 1, 2, or 3 ^2^H_7_-iso-C16:0 containing lipids within the indicated lipid class. TAG, triacylglycerol; PC, phosphatidylcholine; PE, phosphatidylethanolamine, SM, sphingomyelin.

**Table 2.** Table summarizing the acyl chain composition of each [M+NH_4_]^+^ TAG 48:0 with 0, 1, 2, or 3 ^2^H_7_-iso-C16:0 acyl chains incorporated. The percent composition of each acyl chain of the TAG was assessed by comparing the MS2 spectra for each peak. Calculated percentages are comparable to the theoretical composition of the specific acyl chain of interest: fully incorporated (100%), one-third (33.3%) incorporated, or two thirds (66.7%) incorporated.

| Lipid | RT (min) | Formula | Exact Mass | Precursor Ion (m/z)<br>[M+NH<sub>4</sub>]<sup>+</sup> | Diacyl Product Ion<br>(m/z) | Fatty Acyl Chain<br>Loss | % Acyl Chain<br>Composition |
| --- | --- | --- | --- | --- | --- | --- | --- |
| TAG (48:0) | 26.32 | C<sub>51</sub>H<sub>98</sub>O<sub>6</sub> | 806.7363 | 824.7702 | 551.0519 | C16:0 | 100% |
| TAG (48:0)<br>1 <sup>2</sup>H<sub>7</sub>-iso-C16:0 | 26.09 | C<sub>51</sub>H<sub>91</sub>D<sub>7</sub>O<sub>6</sub> | 813.7797 | 831.8141 | 551.0519<br>558.5460 | <sup>2</sup>H<sub>7</sub>-iso-C16:0<br>C16:0 | 34.02%<br>65.98% |
| TAG (48:0)<br>2 <sup>2</sup>H<sub>7</sub>-iso-C16:0 | 25.89 | C<sub>51</sub>H<sub>84</sub>D<sub>14</sub>O<sub>6</sub> | 820.8237 | 838.8560 | 558.5460<br>565.5892 | <sup>2</sup>H<sub>7</sub>-iso-C16:0<br>C16:0 | 65.16%<br>34.84% |
| TAG (48:0)<br>3 <sup>2</sup>H<sub>7</sub>-iso-C16:0 | 25.66 | C<sub>51</sub>H<sub>77</sub>D<sub>21</sub>O<sub>6</sub> | 827.8676 | 845.9020 | 565.5892 | <sup>2</sup>H<sub>7</sub>-iso-C16:0 | 100% |

Next, we quantified incorporation of ^2^H_7_-iso-C16:0 across different lipid classes within the treated 3T3-L1 adipocyte cultures. We observed our tracer incorporated into one or both acyl chains for phosphatidylcholine (PC) species such as PC 32:0 and PC 34:0 (Fig. 3B and Supp. Fig. 3B-C). Interestingly, when analyzing compound lipid species with unsaturated acyl chains we did not see full incorporation of our tracer. For example, for TAG 48:1 we could only observe up to two ^2^H_7_-iso-C16:0-derived acyl groups, and for PC 34:1, only one acyl chain was our incorporated tracer. This data suggests that the iso-C16:0 is not desaturated by 3T3-L1 adipocytes.

We also detected significant ^2^H_7_-iso-C16:0 incorporation into PE 34:0 and ether lipids such as PC O-32:0 (as previously observed(5), Fig. 3B and Supplemental Fig. S3E-F). In the sphingolipid pathway, we observed low but quantifiable incorporation into SM 34:1 and Ceramide d18:1/16:0 species (Fig. 3B and Supplemental Fig. S3G-H). In our hands, differentiated 3T3-L1 adipocytes do not exhibit high rates of sphingolipid biosynthesis, relying more on neutral lipid synthesis for lipid droplet formation. Presumably, cells with a high rate of sphingolipid biosynthesis (e.g. epithelial cells) will exhibit a higher propensity to incorporate BCFAs into sphingoid bases and ceramides(27). Regardless, our integrated results leveraging uniquely labeled tracers and Bckdha-deficient 3T3-L1 adipocytes highlights the ability of C30 chromatography to resolve branched-chain acyl diversity in compound lipids.

### Amino acid tracing of synthesized BCFA into compound lipids

Finally, we additionally confirmed the presence of BCFAs in TAG species by labeling amino acid precursors to these lipids so that we could detect their endogenous production and incorporation into compound lipids. We specifically chose [U-^13^C_5_]valine for these studies, as its catabolism yields labeled isobutyryl-CoA (which is elongated to form iso-C16:0) or propionyl-CoA (which is elongated to form OCFAs). As such, iso-C16:0 produced from [U-^13^C_5_]valine will show M+4 labeling and OCFAs will show M+3 mass shifts (Supplemental Fig. 4). 3T3-L1 adipocytes were cultured in unlabeled or labeled valine and with or without the addition of B12 supplementation, which promotes propionyl-CoA entry to the TCA cycle via methyl-malonyl-CoA mutase and OCFA levels. Exemplary TAG species of 48:0 and 47:0 are discussed in detail from these data. Straight-chain TAG 48:0 exhibited little to no isotopic labeling from [U-^13^C_5_]valine (Fig. 4A-right). In contrast, the smaller BCFA containing TAG 48:0 showed ~60% M+4 labeling, confirming the presence of an iso-C16:0 group and highlighting the significant turnover of the branched-TAG 48:0 pool (Fig. 4A-left). Notably, the addition of vitamin B12 to cultures caused a decrease in branched-TG 48:0 and the ratio of branched/straight-chain TG 48:0 (Fig 4B). Although vitamin B12 has not been shown to increase total intracellular iso-C16:0 fatty acid levels, increased flux through the valine catabolic pathway could lead to higher turnover and thus higher labeling of branched-TAG 48:0. In fact, we see lower levels of branched-TAG 48:0 in the +B12 condition (Fig. 4B).

**Figure 4.**
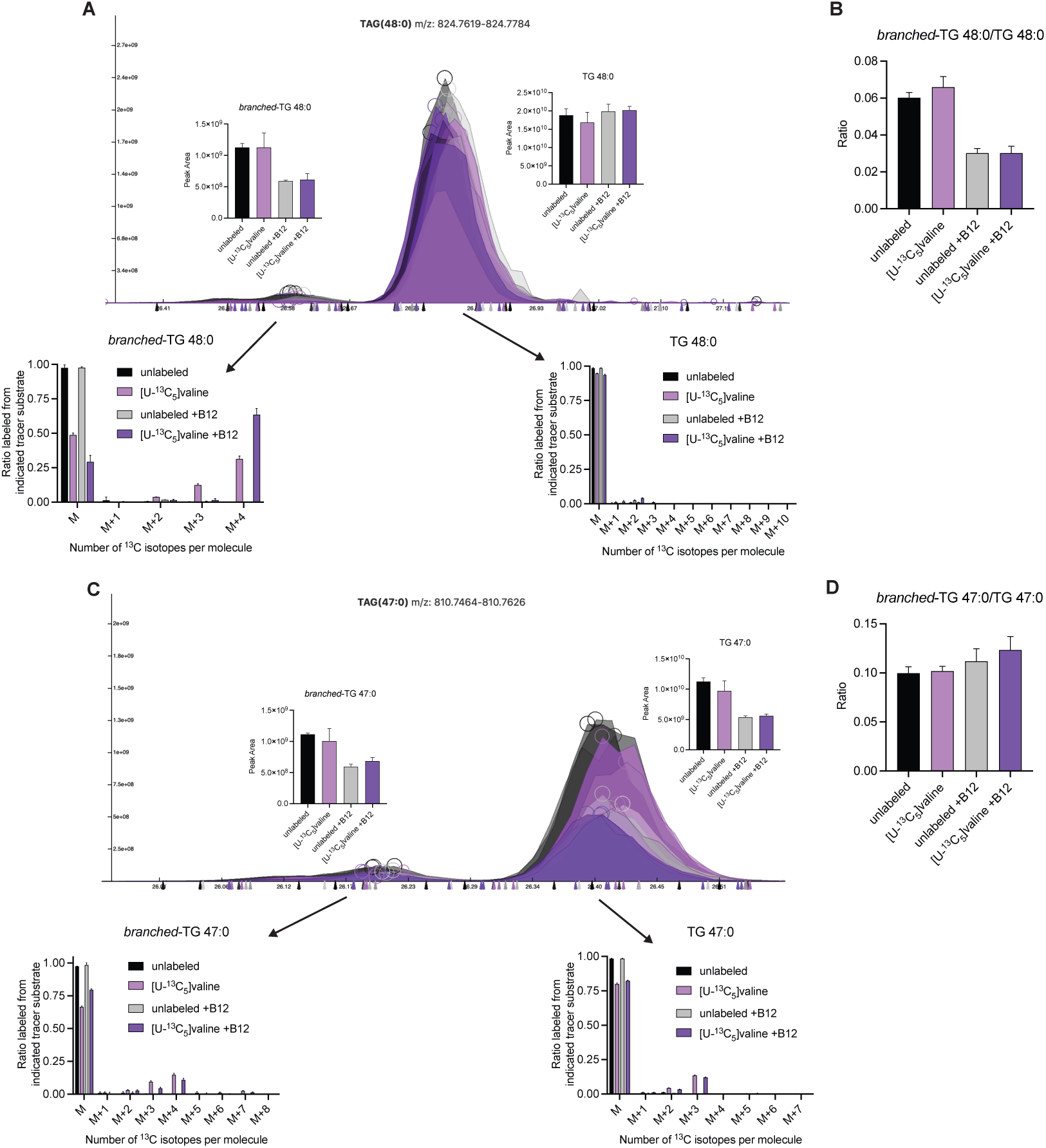
13C-BCAA tracing confirms BCFA incorporation into TAGs. (A) Ion chromatogram of TAG 48:0 traced with [U-13C5]valine. Integrated peak area is depicted within the image and mass isotopomer distribution is below each peak (B) Ratio of branched-TAG 48:0 to straight-chain TAG 48:0 in unlabeled or [U-13C5]valine conditions in the presence or absence of vitamin B12. (C) Ion chromatogram of TAG 47:0 traced with [U-13C5]valine. Integrated peak area is depicted within the image and mass isotopomer distribution is below each peak (D) Ratio of branched-TAG 47:0 to straight-chain TAG 47:0 in unlabeled or [U-13C5]valine conditions in the presence or absence of vitamin B12.

TAG 47:0 also showed putative straight-chain and branched-chain peaks. The larger straight-chain peak showed predominant M+3 labeling indicative of OCFA incorporation (Fig 4C). Interestingly, branched-TAG 47:0 contained both M+3 and M+4 isotopologues (as well as a marginal amount of M+7), providing evidence that this branched-TAG 47:0 peak included both a valine-derived iso-C16:0 and OCFA as acyl-chains (Fig. 4C). Notably, vitamin B12 reduced the abundances of both straight-chain and branched-chain TAG 47:0 peaks, as expected given the presence of OCFAs in both peaks (Fig. 4D). Collectively, the engineering, tracing, and analysis of these adipocytes using C30 chromatography highlights the ability of this approach to deconvolute compound lipids containing branched-versus straight-chain acyl groups in complex biological mixtures.

## Discussion

Here we demonstrate resolution of BCFA-containing compound lipids using an Accucore C30 reversed phase HPLC column coupled to high-resolution mass spectrometry. This column has effectively been used to separate a variety of lipid isomers(17–19). Although GC-MS is effective for separation of methyl esters, recent studies have demonstrated effective resolution of BCFAs as free-fatty acids using reverse-phase liquid chromatography(25). Some studies have also separated BCFA-containing lipids, including lysophospholipids, using tandem mass spectrometric (MS*^n^*) methods via collision-induced dissociation (CID) in human plasma(28)TAGs from *Rhodococcus erythropolis* using RP-HPLC and atmospheric pressure chemical ionization mass spectrometry(29) and diacylglycerols (DAGs) from gram-positive bacteria(30). Still, fractionation of lipids by thin layer chromatography (TLC) followed by GC-MS analysis remains an effective approach to quantify BCFA incorporation into different lipid classes. For example, Liu et al. demonstrated that human fetal intestinal epithelial cells incorporate BCFAs into phospholipids, TAGs, and cholesterol esters(31), similar to our observations with murine 3T3-L1 adipocytes. We used a chemically-synthesized branched fatty acid in addition to metabolically engineered adipocytes, which enabled us to “visualize” the production and incorporation of BCFAs into compound lipids via their distinct labeling patterns. Such approaches are useful for analysis of complex, biologically-synthesized analytes will continue to be valuable tools for exploring new pathways.

The biological function(s) of BCFAs are still being elucidated in higher animals, but in lower organisms they show higher abundance and can influence neuronal development and foraging behavior(32, 33). Recent studies have demonstrated that BCFA-containing lipids exhibit higher affinity for immune receptors. Alpha-galactosyl ceramides containing branched-sphingoid bases exhibit preferential binding to CD1d and signaling through NK cells compared to straight-chain ceramides(34). Alternatively, BCFA-containing phospholipids present in microbes can signal through toll-like receptor 2 (TLR2) to modulate immune cell function(35). BCFAs are consumed in the diet, and levels change in response to dietary fat content. However, BCFAs and OCFAs are highly synthesized and catabolized in mammals from branched-chain CoAs and propionyl-CoA, respectively. The presence of a methyl branch alters the biophysical properties of membranes(36, 37). For example, BCFAs can substitute for polyunsaturated fatty acids to maintain mitochondrial membrane fluidity(38). Accumulation of monomethyl branched fatty acids has shown to be cytotoxic on various cancer cell lines(39–41).

While mass spectrometry can detect the presence of BCFAs in complex lipids, it cannot determine their specific stereochemistry. For instance, in triacylglycerols (TAGs), which contain three stereocenters, identifying the exact location of an incorporated BCFA remains beyond current mass spectrometric capabilities. Furthermore, it is noteworthy that different cell types may exhibit distinct profiles of BCFAs. For example, FADS2, capable of Δ6-desaturating most BCFAs, may not be expressed or has minimal activity in 3T3-L1 adipocytes (42). Moreover, the recently identified enzymes responsible for BCFA elongation could significantly influence the diversity of BCFAs within distinct tissues (43).

In summary, we demonstrate the chromatographic resolution of compound lipids containing monomethyl BCFAs via the utilization of a C30 column. Moreover, employing a metabolically engineered cell line coupled with metabolic tracing techniques confirmed the structures of these complex lipids. The identification and quantification of BCFAs within these intricate lipid compositions offer valuable insights into the metabolic and immunological profiles of patients, holding significant clinical relevance.

## Acknowledgements

We would like to acknowledge past and current members of the Metallo lab for helpful discussions. Support for these studies was provided to C.M.M. by the National Institutes of Health (R01CA234245) and the Lowy Medical Research Institute. Biorender was used to create some Figure graphics.

## Author contributions

C.R.G., M.W., and C.M.M. conceptualization. C.R.G., M.J.K., and G.H.M. formal analysis and investigation. A.T.N. methodology and resources. C.R.G. and C.M.M. Writing – original draft. M.J.K. Writing – review and editing.

## Methods

### Cell Culture & Differentiation

All reagents were purchased from Sigma-Aldrich unless otherwise noted. All media and sera were purchased from Life Technologies unless otherwise stated. Mouse 3T3-L1 pre-adipocytes were generously gifted to our lab by Alan Saltiel and cultured in high glucose Dulbecco’s modified Eagle medium (DMEM) supplemented with 10% bovine calf serum (BCS) below 70% confluence. Cells were regularly screened for mycoplasma contamination. For differentiation, 10,000 cells/well were seeded onto 12-well plates and allowed to reach confluence (termed Day-1). On Day 0, differentiation was induced with 0.5 mM 3-isobutyl-1-methylxanthine (IBMX), 0.25 μM dexamethasone, and 1 μg/ml insulin in DMEM containing 10% FBS. Medium was changed on Day 3 to DMEM + 10% FBS with 1 μg/ml insulin. Day 6, and thereafter, DMEM + 10% FBS was used. Cobalamin (500 nM) was supplemented to cultures when noted. [U-^13^C_5_] valine tracing began 5d post-induction of differentiation. Cells were incubated in custom DMEM (Hyclone) containing [U-^13^C_5_]valine instead of ^12^C-valine. The cells were incubated in this medium for 96 hours total with a media replacement occurring at 48 hours. [^2^H_7_]-isoC16:0 was obtained from LifeLipids, LLC and confirmed for specificity via GC-MS. FA tracing and all FA treatments began 6d post-induction of differentiation. Cells were incubated in DMEM + 10% delipidated FBS + 100μM BSA-conjugated FA or FA tracer. Briefly, this was prepared through preparation of a 100mM FA stock solution in ethanol. This was added in a 1:50 ratio to 4.4% FA-free BSA in PBS. This yields a well-tolerated 3:1 ratio of FA:BSA. This is used at 5% v/v in cell culture media to produce 100μM treatment(44). Fatty acids were then either calculated as percent total fatty acids and normalized to control conditions or normalized to [^2^H_31_]Palmitate internal standard and normalized to control conditions.

### Extraction of metabolites for GC-MS analysis

For cell culture, polar metabolites and fatty acids were extracted using methanol/water/chloroform with [^2^H_31_]palmitate and norvaline as lipid and polar internal standards, respectively, and analyzed as previously described(45). Briefly, cells were washed twice with saline, quenched with −80C methanol and 4C water containing norvaline, scraped into Eppendorfs, and extracted with chloroform containing [^2^H_31_]Palmitate. After centrifugation, phases are dried separately. Samples were stored at −20 °C before analysis by GC-MS.

### Extraction of lipids for LC-MS/MS analysis

Lipid extraction was carried out using a modified Folch methanol/chloroform/water extraction at a ratio of 5:5:2 with the inclusion of 10nmol C12:0 dodecylglycerol, 10nmol [^2^H_31_]palmitate, 10ng of each lipid in the Avanti EquiSPLASH mix. 2 wells of a 12-well plate were combined to form 1 sample. The methanol phase was washed a second time with chloroform after addition of 2μ formic acid. The chloroform phase (bottom layer) was transferred and dried under nitrogen gas at room temperature. Samples were stored at −20 °C before analysis by LC-MS/MS.

### GC-MS analysis

The dried lower chloroform phase was derivatized to form fatty acid methyl esters (FAMEs) via addition of 500μL 2% H_2_SO_4_ in MeOH and incubation at 50°C for 2h. FAMEs were extracted via addition of 100 μL saturated salt solution and 2 500 μL hexane washes. These were analyzed using a Select FAME column (100m × 0.25mm i.d.) installed in an Agilent 7890A GC interfaced with an Agilent 5975C MS using the following temperature program: 80°C initial, increase by 20°C/min to 170°C, increase by 1°C/min to 204°C, then 20°C/min to 250°C and hold for 10 min.

### LC-MS/MS QQQ and QE analysis

A QExactive orbitrap mass spectrometer with a Vanquish Flex Binary UHPLC system (Thermo Scientific) was used with an Accucore C30 coulmn (2.6 µm, 250 x 2.1 mm, Thermo) at 40°C. 5 μL of sample was injected. Chromatography was performed using a gradient of 40:60 v/v water:acetonitrile with 10 mM ammonium formate and 0.1% formic acid (mobile phase A) and 10:90:0.1 v/v/v acetonitrile:2-propanol:water with 10 mM ammonium formate and 0.1% formic acid (mobile phase B), at a flow rate of 0.2 mL/min. The LC gradient ran from 30% to 43% B from 3-8 min, then from 43% to 50% B from 8-9 min, then 50 to 90% B from 9-18 min, then 90 to 99% B from 18-26 min, then held at 99% B from 26-30 min, before returning to 30% B in 6 min and held for a further 4 min.(17) For C18 chromatography studies we used a Kinetex C18 column (1.7 µm, 250 x 2.1 mm, Phenomenex) with the same parameters as stated above.

TAGs, phospholipids, and sphingolipids were analyzed in positive mode using spray voltage 3.2 kV. Sweep gas flow was 1 arbitrary units, auxiliary gas flow 2 arbitrary units and sheath gas flow 40 arbitrary units, with a capillary temperature of 325°C. Full MS (scan range 200-2000 m/z) was used at 70,000 resolution with 1e6 automatic gain control and a maximum injection time of 100 ms. Data dependent MS2 (Top 6) mode at 17,500 resolution with automatic gain control set at 1e5 with a maximum injection time of 50 ms was used. Extracted ion chromatograms for each analyzed lipid was generated using a m/z ± 5 ppm mass window around the calculated exact mass. Peak areas for TAGs were normalized to Avanti EquiSPLASH internal standard (specifically TAG 15:0-18:1(d7)-15:0).

### Statistical analyses

All experiments were repeated at least 2 times and data from 1 representative experiment is shown. For analyses involving 2 groups, a two-tailed Student’s t-test was performed. For analyses with 3 groups or more, a one- or two-way ANOVA was performed as appropriate. Corrections for multiple comparisons were performed for all analyses except Seahorse measurements.

---

**Supplemental Figure 1.**
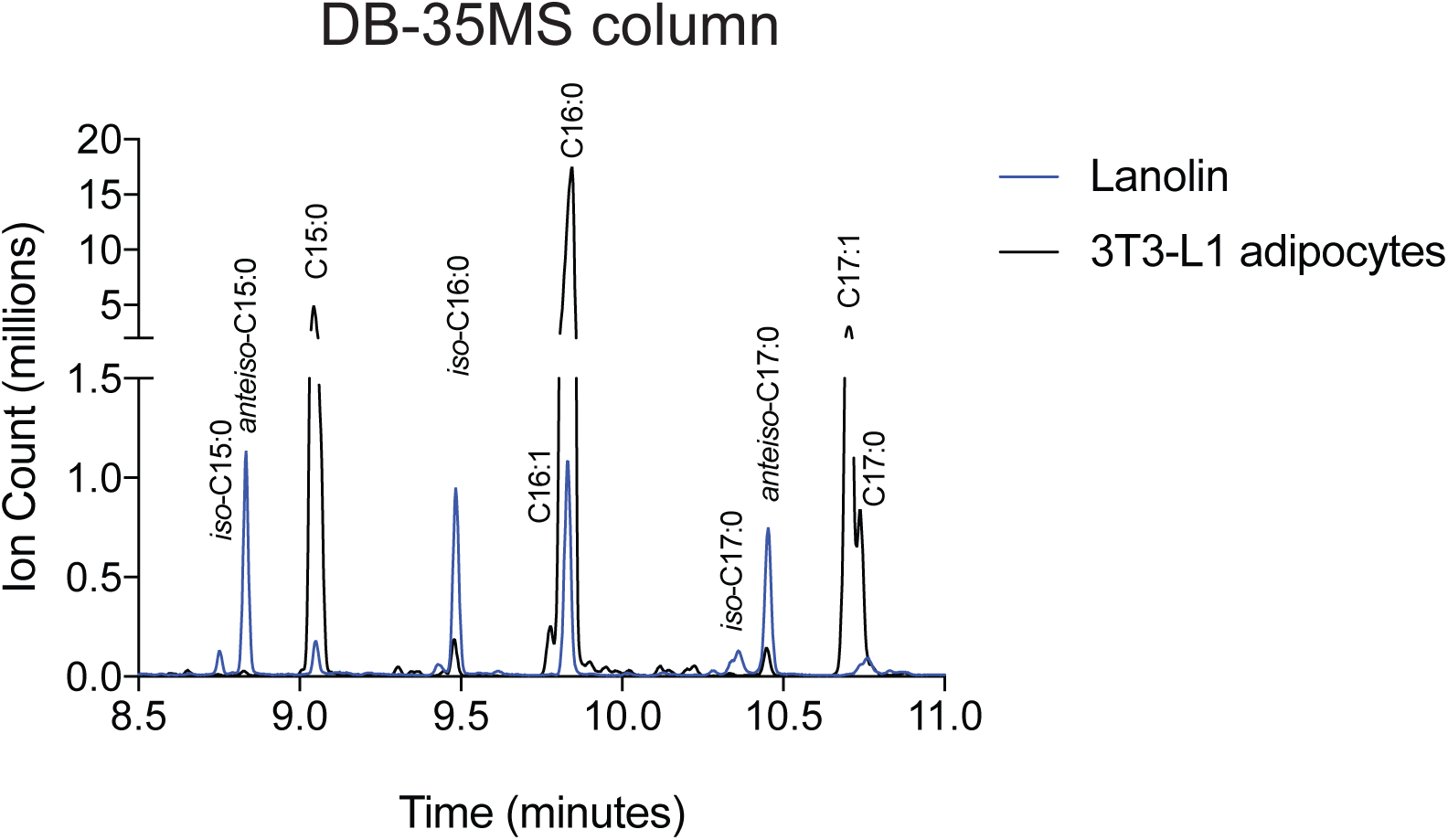
Branched- and straight-chain fatty acids are chromatographically resolvable on GC-MS. Separation of free monomethyl BCFAs and straight chain fatty acids as fatty acid methyl esters in lanolin extract and 3T3-L1 adipocytes using via GC-MS using a DB-35MS column.

**Supplemental Figure 2:**
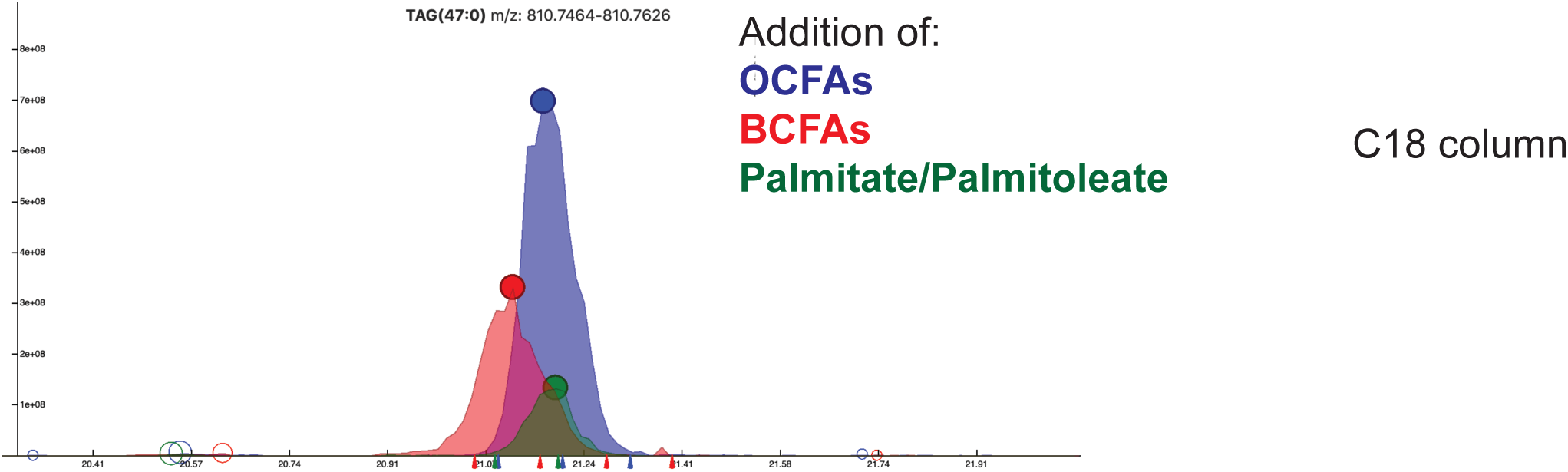
Experimental conditions from **Figure 1**, ran on Kinetex C18 column (1.7 µm, 250 x 2.1 mm, Phenomenex).

**Supplemental Figure 3:**
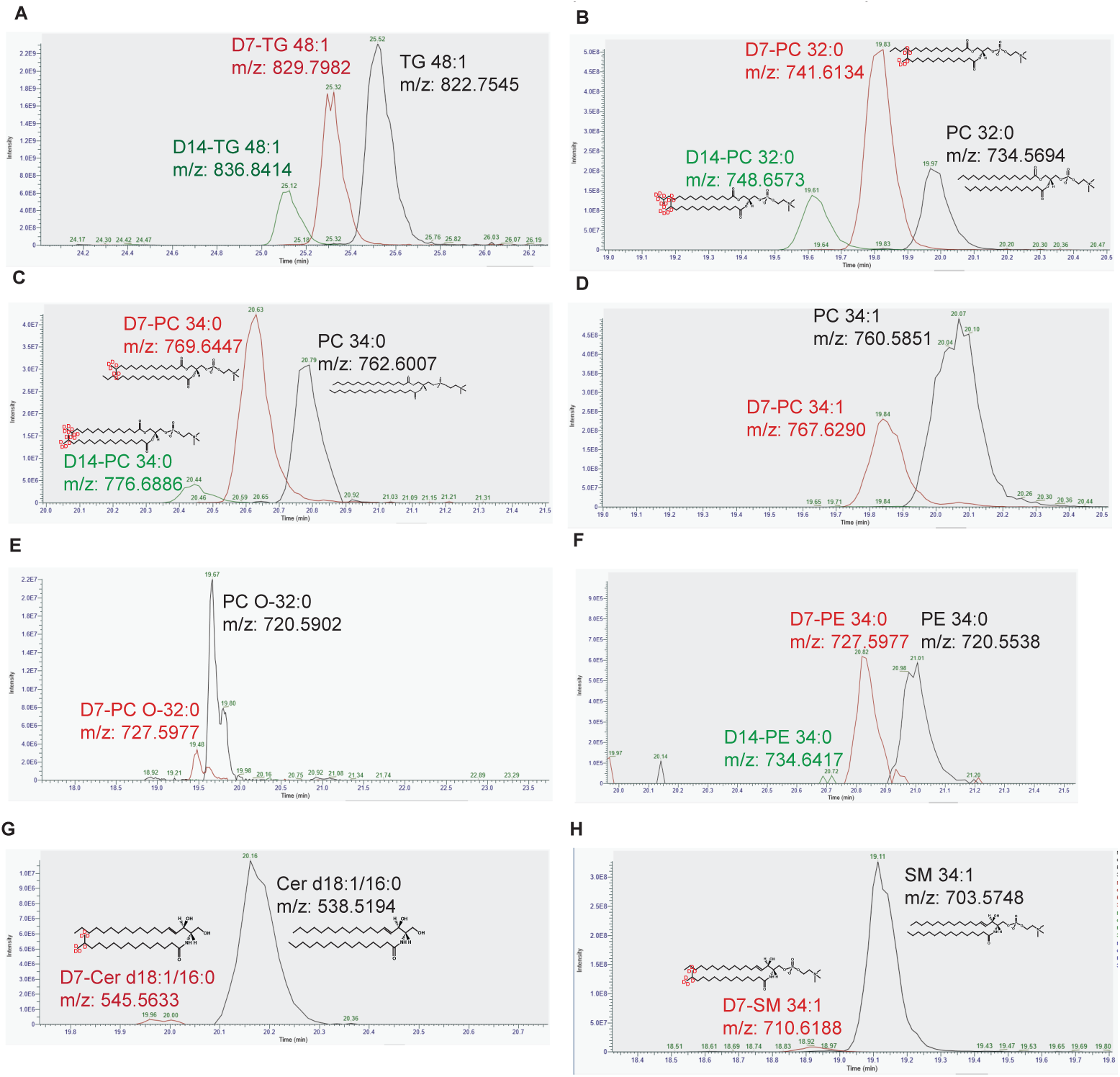
(**A-H**) Extraction ion chromatograms illustrating ^2^H_7_-Iso-C16:0 integration across different complex lipids. Retention time shifts highlighted with the incorporation of each BCFA acyl chain. Respective extracted ion *m/z* displayed in each figure; ion adduct extracted shown below for each class of lipids. TAG, triacylglycerol; PC, phosphatidylcholine; PE, phosphatidylethanolamine, SM, sphingomyelin. (A) TAG 48:1 [M+NH_4_]^+^ (B) PC 32:0 [M+H]^+^ (C) PC 34:0 [M+H]^+^ (D) PC 34:1 [M+H]^+^ (E) PC O-32:0 [M+H]^+^ (F) PE 34:0 [M+H]^+^ (G) Ceramide d18:1/16:0 [M+H]^+^ (H) SM 34:1 [M+H]^+^

**Supplemental Figure 4:**
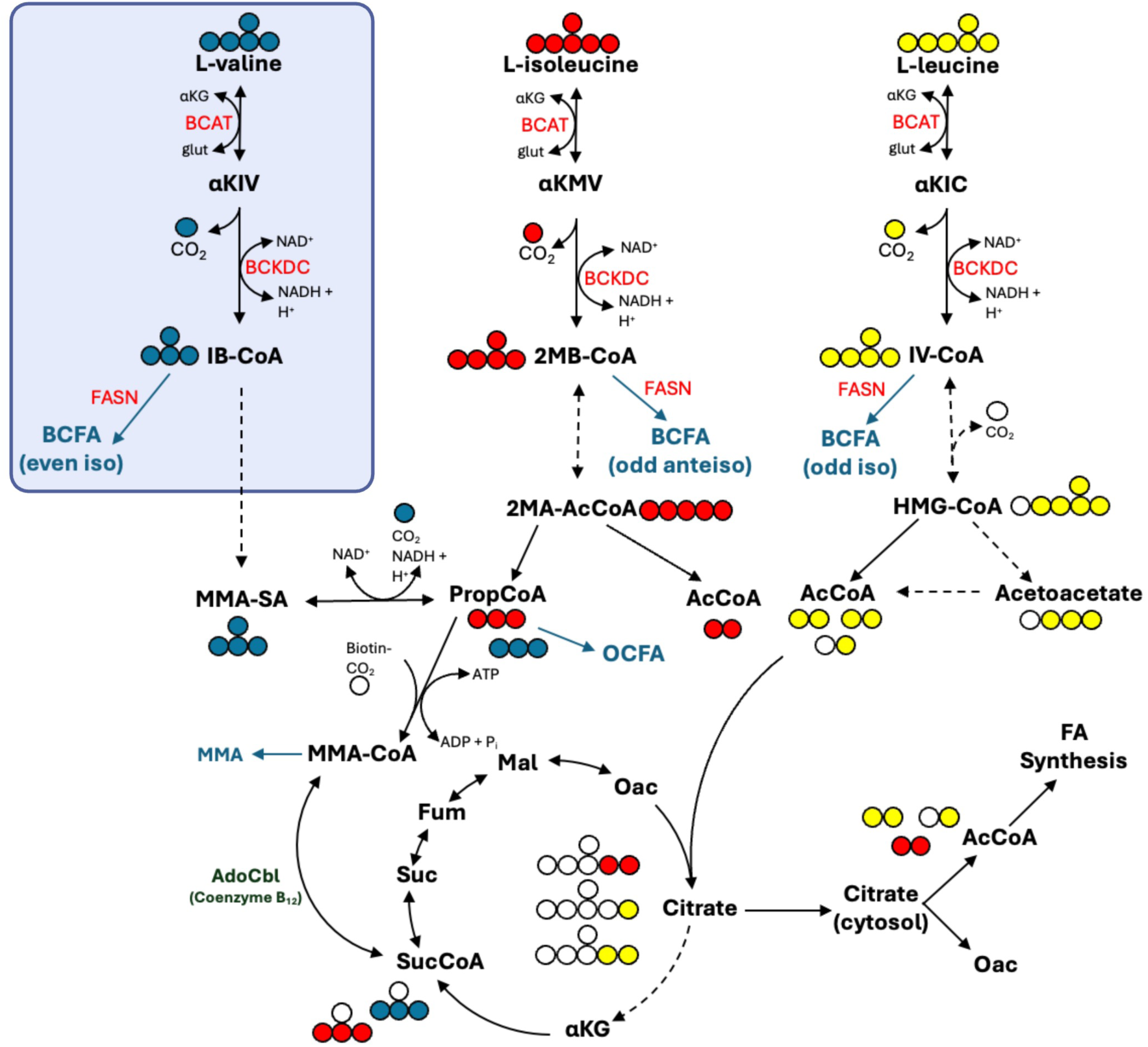
Metabolic map depicting the catabolism of the branched chain amino acids (BCAA) valine, isoleucine, and leucine. Oxidation of [U-13C5]valine leads to BCFA synthesis (highlighted in the blue box), OCFA synthesis, and TCA cycle incorporation. Key enzymes for BCFA biosynthesis highlighted in red text. BCAT, branched-chain amino acid aminotransferase; BCKDC, branched-chain a-ketoacid dehydrogenase complex; FASN, fatty acid synthase.

